# Environmental Toxin Rotenone Drives LRRK2-Mediated Microtubule, Cilia and Proteostasis Disruption in Parkinson’s Disease Model

**DOI:** 10.64898/2026.01.08.697160

**Authors:** Dani Flinkman, Prasannakumar Deshpande, Peter James, Eleanor Coffey

## Abstract

Exposure to the environmental toxin rotenone increases risk for Parkinson’s disease (PD). However, protein phosphorylation changes induced by rotenone in neural-derived cells have not been reported. We examined the effect of rotenone on the proteome and phosphoproteome of cortex-derived cultures from wild-type and *Lrrk2-/-* mouse brains. We also analyzed the phosphoproteome of the PD cadaver brain using a previously unanalyzed dataset. Rotenone alters phosphorylation at 904 sites, most of which are unchanged in *Lrrk2-/-* cultures. Common targets include proteins that control microtubule stability, vesicular transport, and protein clearance via the autophagosomal/lysosomal pathway. Analysis of phosphosites with known function indicate that rotenone activates histone deacetylase-1 and represses the pro-survival transcriptional regulators Mef2C and Mef2D, while inhibiting translation by phosphorylating Eif2b5. These effects do not occur in *Lrrk2-/-* cultures. This study shows that *Lrrk2* is required the earliest rotenone-triggered phospho-signalling events.

## INTRODUCTION

Parkinson’s disease (PD) develops due to both genetic and environmental factors^1,2^. Exposure to insecticides like rotenone, and the herbicide paraquat, is known to raise the to develop PD^3,4^. In rat models, rotenone induces selective loss of substantia nigra (SN) dopaminergic neurons and causes *α*-synuclein aggregation, reminiscent of Lewy bodies. It also induces motor deficits and non-motor symptoms such as hyposmia and cognitive decline^5,6^.

Rotenone, paraquat and MPP^+^ (cyperquat metabolite) all inhibit tubulin self-assembly^7–9^. Rotenone does this by binding to the colchicine site on microtubules^10^. The highly arborized axons of dopaminergic neurons renders them particularly vulnerable to microtubule disruption^11,12^. Rotenone induces microtubule depolymerization and axonal shrinkage in midbrain neurons^13,14^, which may explain its effects in PD. Rotenone also inhibits mitochondrial complex I, leading to oxidative stress^15^. Interestingly, rotenone activates Leucine-rich repeat kinase 2 (LRRK2), which is linked to both familial and sporadic forms of PD^13,16^. LRRK2 oligomerizes on microtubules promoting their stability^17^. Blocking LRRK2 can protect neurons from rotenone toxicity, suggesting a causal link^13,18,19^. This study utilizes a discovery-driven, shotgun proteomics approach to examine the early effects of rotenone on the proteome and phosphoproteome of cortex-derived cultures from wild-type (WT) and *Lrrk2-/-* mice. The aim is to identify the molecular changes induced by rotenone in neural-derived cultures and to identify mechanisms that require Lrrk2.

## RESULTS AND DISCUSSION

### Ciliary proteins

We profiled protein expression in cortex-derived cells (consisting of mixed neurons and glial cells) from mice lacking Lrrk2 (Fig.1). This revealed a marked regulation of cilia proteins. In all, 118 cilia proteins were significantly altered (FC>1.5, FDR<0.05), 86% of which were upregulated (Fig.1a). The largest changes were for Lrrc46 and Lrrc23, proteins that regulate cilia (and flagella) spoke formation^20^. These proteins are expressed at high levels in the midbrain^21^. Radial spoke head proteins 1 (Rsph1) and 4 (Rsph4a), and ciliary dyneins were also upregulated (Fig.1a-c,f). Most of these cilia proteins shared a GO term with the keyword microtubule. This expansive upregulation of cilia proteins in *Lrrk2-/-* cells is consistent with previous studies showing. Those had shown that the activating LRRK2-G2019S mutation reduces cilia number in the brain^22,23^.

**Figure 1.**
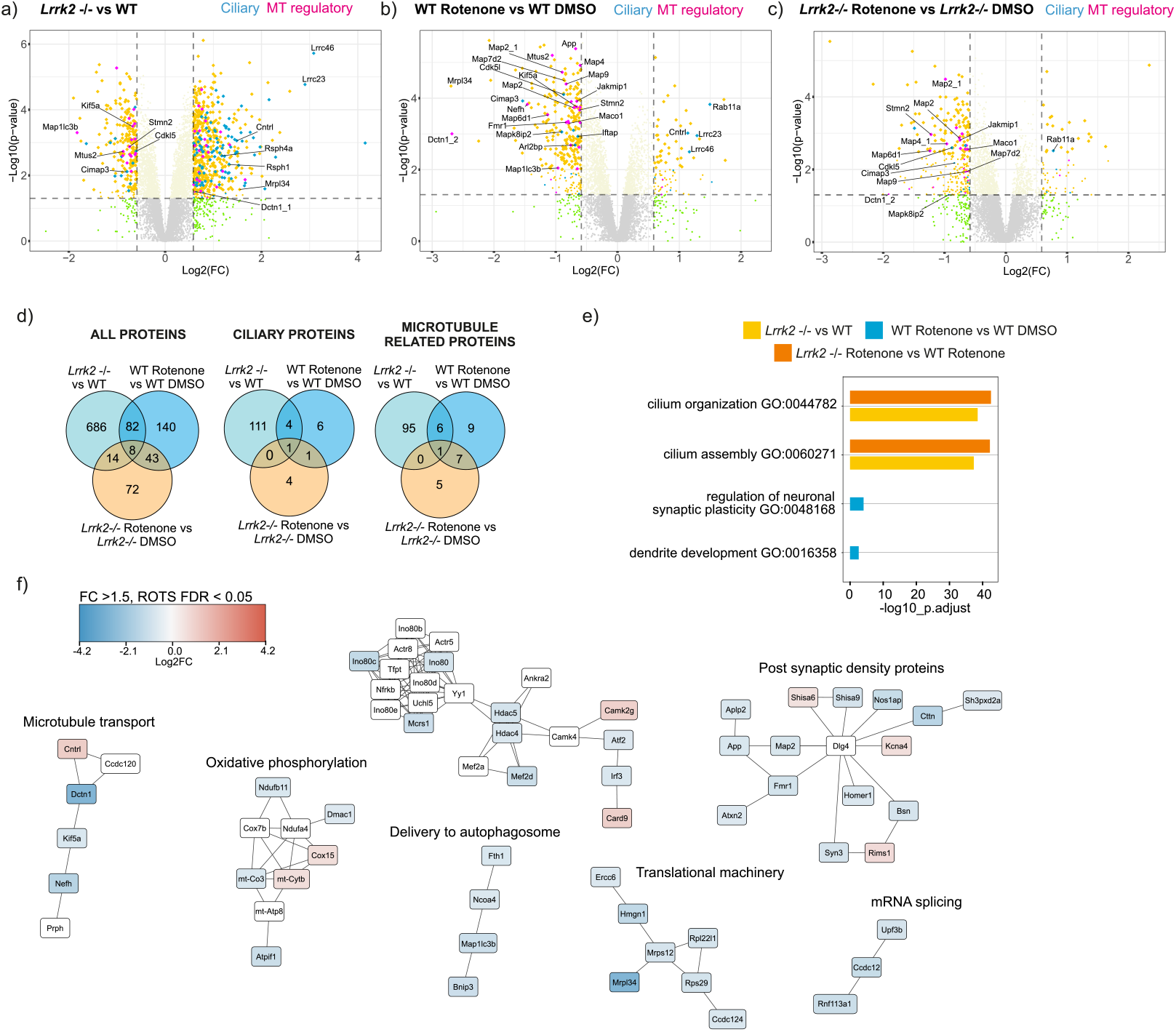
Rotenone-induced protein abundance changes in cortical tissue cultures (5 replicates for *Lrrk2-/-* and 6 for WT) are shown as volcano plots for **a)** *Lrrk2-/-* vs wild-type (WT), **b)** 100 nM rotenone-treatment WT vs WT DMSO, **c)** *Lrrk2-/-* treated with 100 nM rotenone vs *Lrrk2-/-* DMSO with FC>1.5 and p<0.05. Proteins with FDR<0.05 are yellow diamonds, those mapping to Cilia GO-terms are turquoise, and to microtubule GO-terms are magenta. Microtubule proteins mapping to cilia are manually assigned based on predominant characterized functions. Lrrc24 and 46, lacking GO terms, are mapped to cilia manually. **d)** Venn diagrams show significantly changing proteins with FC>1.5, FDR<0.05 for each comparison including overlap. P-values and FDR were calculated using the Reproducibility Optimized Test Statistic (ROTS). **e)** Gene ontology analysis depicts the top two GO terms for each comparison. **f)** Protein-protein interaction (PPI) network of rotenone-induced protein expression changes: increases (red), decreases (blue), and inferred nearest neighbours not identified (white). For compactness, proteins are referred to by gene names throughout.

We next examined the effect of acute rotenone treatment on protein expression in cortical cultures. Rotenone downregulated ten and upregulated two ciliary proteins (Fig.1b-c,Suppplementary Data 1,2.). Specifically, Dctn1, Cdk5l, App, ADP ribosylation factor 2 binding protein (Arl2bp), Bbip1, ciliary microtubule interacting protein-3 (Cimap3), Kif5A and Iftap decreased, whereas centriolin (Cntrl) and Rab11a increased. Furthermore, known ciliary proteins Lrrc23 and Lrrc46, that did not map to ciliary GO terms were upregulated. Notably, all of these effects (except for Rab11a and Cdk5l) were ablated when rotenone treatment was done in *Lrrk2-/-* cells (Fig. 1c). Overall, the effect of rotenone on ciliary proteins less extensive than *Lrrk2* deletion (Fig. 1a-d).

### Microtubule and protein transport proteins

Besides ciliary proteins, *Lrrk2* and rotenone altered the levels of proteins that control microtubule stability and protein transport (Fig.1a-e). Accordingly, rotenone decreased Map2, Map4, Map9, Nefh, Fmr1, Map1lc3b, Dctn1, Kif5A, Map6d1, Map7d2, Mtus2, Maco1, Jakmip1 and Stmn2 (Fig. 1b). Among these, Stmn2 stood out as a microtubule regulator essential for axonal integrity^24,25^. Moreover, *STMN2* is implicated in PD and in ALS, and *Stmn2* knockdown induces substantia nigra dopaminergic neuron atrophy in vivo^26^. MAP4 also stooud out as we have previously reported that its expression is decreased in fibroblasts from sporadic PD cases^27^.

### Autophagosome-related proteins were also dysregulated by rotenone

LC3B (Map1lc3b), essential for autophagy, mitophagy and mRNA degradation^28^, reduced by 2-fold (FDR=0.04) in rotenone-treated cultures and by -3.5-fold (FDR=0.002) in *Lrrk2-/-* cells, but not in rotenone-treated *Lrrk2-/-* cells (Fg.1a-f). Similarly, dynactin-1 isoform-2 decreased (-6.4-fold, FDR=0.0) in rotenone-treated WT, but not in *Lrrk2-/-* cells. These proteins are part of the dynactin complex that links dynein to lysosomes. Mutations in dynactin are associated with late-onset Parkinsonian disorders such as Perry syndrome^29^. The dynactin binding scaffold Mapk8ip2 (JIP2) also decreased (-1.9-fold, FDR=0.01) following rotenone (Fig.1a-c,f). Besides autophagosome delivery proteins, components of the translational machinery were also downregulated (Fig. 1f). Down regulation of proteins involved in autophagosomal/lysosomal and mitochondrial homeostasis, may induce oxidative stress, inflammation and diminished clearance of protein aggregates^30^.

### Phosphorylation regulation

Given that rotenone activates LRRK2^13,16^, we investigated phosphorylation changes in cortex-derived cultures from WT and *Lrrk2-/-*. This identified 9713 phosphosites. As many did not meet the FDR cut-off of < 0.05 for FC>1.5, we relaxed the significance threshold to p-value<0.05. Rotenone induced 904 phosphosite changes, of which 783 occurred in rotenone-treated WT neurons but not in rotenone-treated *Lrrk2-/-* neurons, suggesting that Lrrk2 expression is required (Fig. 2a-c). These phosphorylation changes were highly enriched for the GO-term “*tubulin binding*” (FDR= 0.001). Functional annotation was available for 77 of these phosphosites, as discussed below.

**Figure 2.**
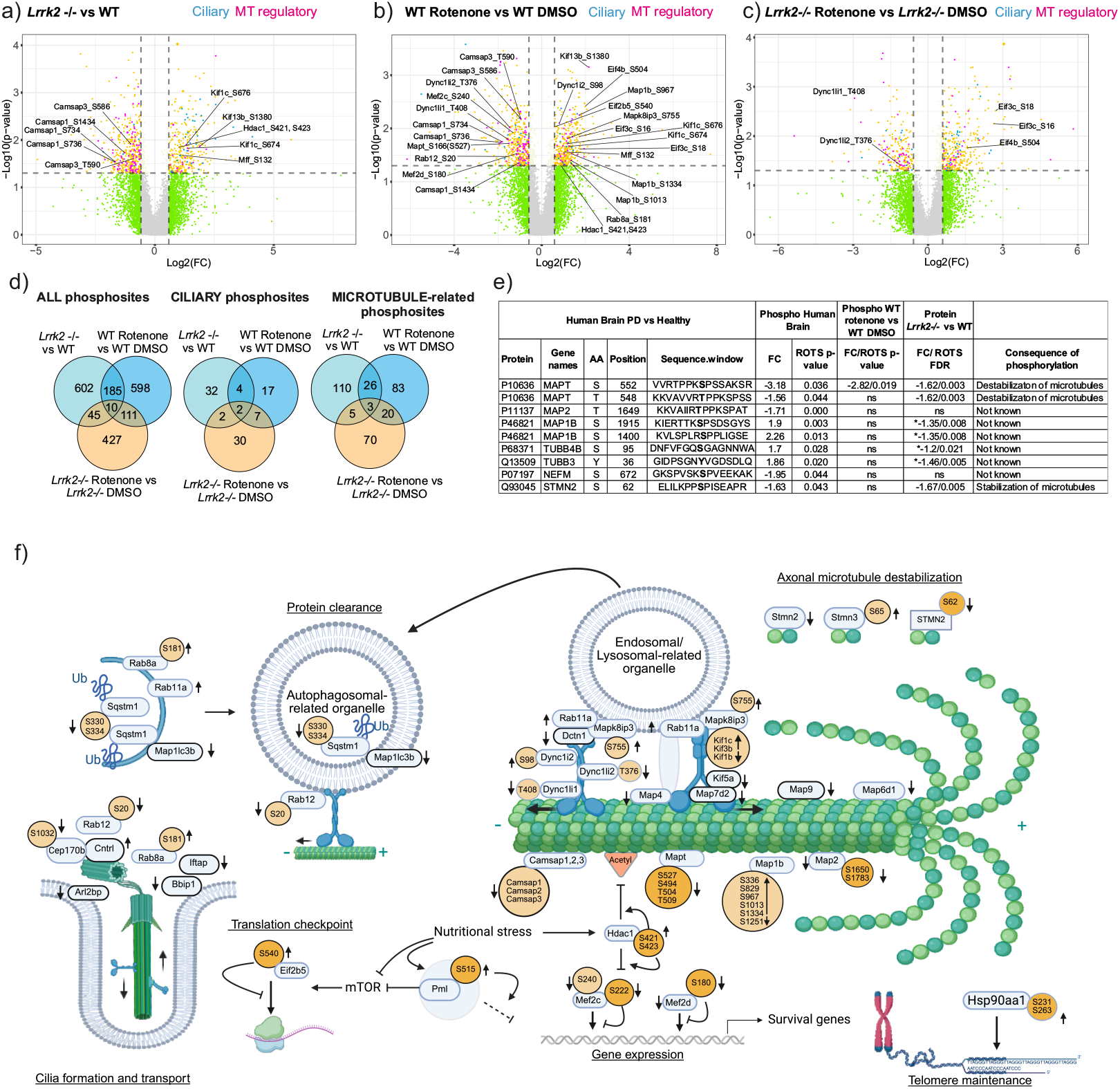
Rotenone-induced changes in cortex-derived tissue cultures (n=6 per condition). **a)** *Lrrk2-/-* vs WT, **b)** rotenone vs DMSO in WT, and **c)** rotenone-treated vs untreated *Lrrk2-/-* cultures. Volcano plots highlight phosphosites (FC>1.5, p<0.05). Phosphosites with FDR<0.05 are diamonds; those on cilia proteins are turquoise, those from microtubules are magenta. Microtubule proteins mapping to cilia are manually assigned based on main functions. **d)** Venn diagrams show overlapping changing phosphosites for indicated groups, FC>1.5, p<0.05. **e)** Phosphorylation changes in temporal gyrus from PD brain (FC>1.5, p<0.05), from 5 female patients and 5 controls aged 81-84. Corresponding cortical culture data is shown (FC>1.5, p<0.05). * indicates FDR<0.05, FC<1.5. **f)** Model depicts proteins and phosphosites regulated by rotenone, contributing to PD mechanism. Phosphosites (yellow) have known function; pale orange are unknown. Fold-change for expression (grey) and phosphorylation (orange) is indicated by up/down arrows. Rotenone-affected proteins/phosphosites found only in WT are outlined in black. The model shows that phosphorylation of proteostasis mechanisms, cilia formation, microtubule cytoskeleton, and protein translation are all regulated, as discussed. The cartoon was created in BioRender.com^48^ https://BioRender.com/wxwqfe5.

### Transcriptional regulator phosphorylation

Among the annotated sites, rotenone decreased phosphorylation of transcription factors Mef2c-S240 (-2.6-fold, p=0.02) and Mef2d-S180 (-2.5-fold, p=0.03) on their trans-activating sites (Fig. 2a-d). Mapk7 (ERK5) phosphorylates these sites in response growth factors and induces pro-survival gene expression^31,32^. This rotenone-induced downregulation was absent in *Lrrk2-/-* neurons. Furthermore, transcriptional regulator Pml-S515 phosphorylation increased 2.5-fold (p=0.001). PML plays multifaceted roles, negatively regulating mTOR and translation initiation. Phosphorylation on S515 impacts its ability to regulate transcription^33^. Chromatin remodelling enzyme histone deacetylase (Hdac1) was also regulated. Specifically, Hdac1-S423 and S421 phosphorylation was increased in rotenone-treated neurons (Fig. 2b), while Hdac1 protein level was unchanged. Phosphorylation on these sites increases its deacetylase activity leading to chromatin compaction and repressed transcription^34^. Moreover, HDAC1 deacetylates tubulin K40 in response to nutritional stress^35^. This can destabilize axonal microtubules (which are highly acetylated) and may contribute to neurodegeneration^36^. Additionally, HDAC1 inhibitors are neuroprotective in PD models^37^. Regulation of Hdac1 phosphorylation does not occur when rotenone is applied to *Lrrk2-/-* cortical cultures, suggesting that LRRK2 is required.

### Protein synthesis regulator phosphorylation

Rotenone increased the phosphorylation of the translational initiation factor Eif2b5 (Eif2b*ϵ*) on several sites including S540 (+1.5FC, p=0.009) (Fig. 2b). Eif2b5-S540 phosphorylation disrupts its ability to recycle Eif2-GTP. This prevents tRNA ternary complex formation and subsequent translation initiation^38^. Eif2b5-S540 phosphorylation is highly studied in the context of the integrated stress response^39^, where phosphorylation of this site represses translation. Mutations in Eif2b5 are associated with various genetic cases of vanishing white matter diseases, demonstrating its critical function in maintaining axonal integrity in particular^40^. Diminished white matter is also observed in the ageing brain and in PD^41^. There was no change in Eif2b5 phosphorylation on any sites in rotenone-treated *Lrrk2-/-* cells. Interestingly, rotenone substantially reduced expression of the mitochondrial translational regulator Mrpl34 (-6.5-fold, FDR=0) (Fig. 1a-g). This is expected to lead to rundown of oxidative phosphorylation. These results suggest that translation is rapidly reduced in response to rotenone. They support earlier reports linking rotenone and LRRK2-dependent repression of translation as an early event in PD^13,42,43^.

### Proteostasis and lysosomal trafficking complex phosphorylation

Rotenone also regulated the phosphorylation of axonal transport machinery. For example, microtubule-associated protein 1b (Map1b)-S967, S1013 and S1334, Mapk8ip3 (JIP3)-S755, dynein intermediate chain isoform 2 (Dynci2)-S98 and Rab8a-S181 all showed a significant phosphorylation increase (Fig. 2a-f). These changes were absent in rotenone-treated *Lrrk2-/-* neurons. While the function of these sites remains unannotated, the proteins are known to direct axonal transport. JIP3 (like JIP4) is an activating adaptor that tethers autophagosomal/lysosomal cargo to the dynein-dynactin complex (linked via Dyncli1 or 2). JIP3 binding promotes “retrograde” cargo transport^44^. This is further stabilized by Map4^45^, which decreases in expression following rotenone (-1.5 FC, FDR=0; Fig1.b), implying that soma-direct transport is repressed. Retrograde transport is further challenged when LRRK2 is activated leading to hyperphosphorylation of Rab proteins e.g. Rab12, Rab8 and Rab10. Phosphorylated Rabs recruit kinesin-1 to the autolysosome, facilitating anterograde transport (Fig. 2g). This in turn reduces clearance of autolysosomes from axons^44,46^. Here we observe that rotenone regulates the phosphoryation of Rab8a-S181 (+2.5 FC, p=0.04) and Rab12-S20 (-3FC, p=0.03), pivotal players in this mechanism. Furthermore, we observed decreased S334/S340-Sqstm1 (p62) phosphorylation following rotenone (Fig.2). This protein sequesters ubiquitin-tagged proteins to the autophagosome, mediated by LC3/ATG8 binding^47^. It plays a major role in retrograde transport of aggregated proteins and is implicated in PD and ALS. Although these sites are not functionally annotated, their regulation suggests that rotenone may regulate autophagosome maturation into an autolysosome and mechanisms of protein clearance in the axon. The consequence may be toxic accumulation of proteins.

To connect these findings to a clinical context, we analysed the MS-DIA spectra from the temporal gyrus of post-mortem PD cases to gain insight on phosphorylation changes in PD brain^49^. Reduced grey matter and volume in this region is associated with declined cognition and language processing in PD^50^. We found reduced STMN2-Ser62 phosphorylation, which destabilizes microtubules and signifies axonal damage^24,26,51^. Several other MAPs, and tubulin isoforms, showed significantly altered phosphorylation (Fig.2e). The same MAPs were altered in mouse cultures following rotenone treatment, but on different residues. One exception was MAPT (Tau)-S552 phosphorylation which was decreased in PD brain and in cortical cultures treated with rotenone. Phosphorylation on this site is highly studied Alzheimer’s disease where it promotes fibril formation^52^. Notably, neither STMN2-S62 nor MAPT-T552 phosphorylation were altered by rotenone in *Lrrk2-/-* cultures.

### Conclusions

This study reports the early effects of rotenone treatment on protein expression and phosphorylation in cortical tissue cultures.

## METHODS

### Tissue culture isolation and rotenone treatment

*Lrrk2-/-* mice (C57BL/6-Lrrk2tm1) and C57BL/6 WT mice were from the Jackson Laboratory. Cortex was isolated from post-natal day 0 mice as previously described^13^. Cortical tissue was dissected in dissection medium (30 mM K2SO4, 81.8 mM Na2SO4, 5.8 mM MgCl2, 1 mM d-glucose, 0.25 mM Hepes pH 7.4, 0.001% Phenol red, and supplemented with 1 *µ*M kynurenic acid) were digested with Papain (10 U/mL Worthington Biochemical Corp., Lakewood, NJ, USA, 3119) and plated at 900,000 cells/well on 24 well plates. The resulting mixed culture was maintained in MEM (Sigma-Aldrich, St. Louis, MO, USA) supplemented with 10% fetal calf serum, 2 mM Gln and penicillin 50 U/mL and streptomycin 50 *µ*g/ml. At 48h post-plating, media of cells was supplemented with cytosine arabinoside (2.5 *µ*M final) to prevent further glial cell division. This results in a culture of mixed cortical neurons and glial cells. On day 10, cells were treated with DMSO or 100 nM rotenone for 4 hours and thereafter lysed directly in Laemmli buffer (1% sodium dodecyl sulfate (SDS), 10% glycerol, 2.5% 2-mercaptoethanol, 0.005% bromophenol blue in 31 mM Tris-HCl pH 6.8). Protein yield was quantified using the Pierce 660 nm Protein Assay with ionic detergent compatibility kit (Thermo Fisher Scientific, USA).

### MS sample preparation for total protein measurement

Samples were processed with S-Trap 96-well plate (Protifi LLC, Fairpoint, NY, USA). SDS was added to 5% for 40 *µ*g of each sample. Samples were reduced with 5 mM (tris (2-carbxyethyl)phosphine)-HCl for 30 min at RT, followed by alkylation with 20 mM iodoacetoamide in the dark at RT for 30 min. Following alkylation samples were loaded on a S-Trap plate in 100 mM Tris-HCl pH 7.1 buffer in 90% methanol according to manufacturer’s instructions. Proteins were digested with sequencing grade trypsin (Promega, Madison, WI, USA) 1:50 trypsin to protein ratio 16h at 37°C and eluted according to manufacturer’s instruction and dried with SpeedVac (Thermo Fisher Scientific). Samples were reconstituted in 0.1% formic acid and peptide yield was quantified with Pierce™ Quantitative Fluorometric Peptide Assay.

### Phospho-enrichment

140 *µ*g of lysate was loaded on SDS-PAGE gel and resolved 2 cm into a 12.5% resolving gel. Gels were stained with GelCode™ Blue stain and lanes were excised and cut in to 1×1mm pieces, destained and reduced with 20 mM dithiothreitol for 1h at 56°C and alkylated with 55 mM iodoacetoamide for 1h at room temperature in the dark. Protein was digested with sequencing grade trypsin (Promega) at 1:50 trypsin to protein ratio and incubated at 37°C for 16h. Digestion was inactivated by addition of formic acid to final concentration of 0.2%. Digest was collected and peptides were further eluted with acetonitrile (ACN) from the gel pieces. Peptides were dried to full dryness with a SpeedVac (Thermo Fisher Scientific) and enriched with High Select™ TiO2 phosphopeptide enrichment kit (Thermo Fisher Scientific) according to manufacturer’s instructions. Following enrichment samples were dried with a SpeedVac to total dryness and resuspended in 0.1% FA for MS analysis.

### MS analysis

800 ng per total protein samples was loaded to Evotip PURE disposable desalting and trap columns (Evosep Biosystems, Odense, Denmark). Evotips were loaded to Evosep one HPLC (Evosep Biosystems) interfaced with Orbitrap Fusion Lumos (Thermo Fisher Scientific) with FAIMS-Pro (Thermo Fisher Scientific). Evosep one was operated at 30 samples per day mode, FAIMS-Pro with -45V compensation voltage and carrier gas flow rate at 3.5 l/min. Orbitrap Fusion Lumos was operated in data independent acquisition (DIA) mode with the radio frequency (RF) lens set to 35%. MS1 and MS2 scans were obtained with the Orbitrap analyzer with where MS1 scan was obtained at resolution of 120k followed, by 32 DIA scans at 30k resolution at high-energy collisional dissociation (HCD) fragmentation mode with 28% HCD collision energy from mass-range of 449-800 m/z. Normalized automatic gain control (AGC) was set to 125% in MS1 and 800% for MS2 and maximum injection times at 50 ms and 54 ms. For phospho-enriched peptides, the Orbitrap Fusion Lumos was operated in data dependent acquisition mode with RF lense set to 30%. The mass-spectrometer was interfaced with EasynanoLC and peptides were separated in 60 min gradient with C18 column. Peptides were trapped on a 100 *µ*M × 2 cm trapping column and followed by separation on a 75 *µ*M ID × 15 cm analytical column, packed with ReproSil-Pur 5 *µ*m 200 Å C18-AQ (Dr Maisch HPLC GmbH, Ammerbuch-Entringen, Germany) with 300 nl/min flowrate. Buffer A was 0.1% FA and buffer B 80% CAN. In the chromatographic separation buffer B was increased from 5 to 23 min over 29 minutes, followed by increase to 31% from 29 to 50 min, and was increased to 100% in from 50 to 55min and maintained at that until 60min. The mass-spectrometer was operated in 3s cycle mode where both MS1 and MS2 scans were acquired in Orbitrap, where MS1 scan at 120k resolution was acquired every 3s and MS2 scans at electron-tranfer HCD mode were performed between MS1 scans at 30k resolution, and dynamic exclusion was set to 35s. Normalized AGC was set to 100% for both MS1 and MS2 scans, and maximum injection times were set to 50ms and 54ms.

### MS Data analysis

DIA protein data was analysed with Spectronaut 19 (Biognosys AG, Zurich, Switzerland) with sample median normalization and maxLFQ protein summarization turned on. Human brain data was downloaded from PRIDE repository ((PXD022297)^53^. Brain data DIA search was performed as above, but with phosphorylation at serine, threonine and tyrosine as variable modification turned on. MaxQuant 2.6.2.0^54^ was used to analyse phosphorylation data with match between runs turned on. In both cases Mus musculus FASTA from Uniprot was used for database search. In both analyses carbamidomethylation of cysteines was set as fixed modification, and oxidation of methionine and protein n-terminal acetylation were set as variable modifications. Phosphorylation of serine, threonine and tyrosine was set as variable modification for the MaxQuant analysis. Both datasets were pre-processed in Perseus, where proteins and phosphosites, which did not have at least 50% non-missing values in at least one sample group were removed. Phosphorylation data was median normalized and missing values in both datasets were imputed with “Replace missing values from normal distribution” function in Perseus with total matrix selected.

### Statistics and functional data analysis

Statistical analysis was performed using PhosPiR^55^, where ROTS was selected for for p-value and multiple testing correction. GO analysis was performed using org.Mm.eg.db and clusterProfiler packages with Benjamini-Hochberg correction for multiple testing against full mouse genome. Plots were generated using ggplot2. String analysis weas performed with Cytoscape stringApp. In brief confidence score of 0.8 was used for interactions and 30 neighbouring interactors were added to the data with 0.8 interaction confidence score.

## Data availability

The mass-spectrometry raw files from Orbitrap Fusion Lumos have been deposited to the ProteomeXchange Consortium via the PRIDE^56^ partner repository. The data will be made public upon publishing.

## Acknowledgements

Biocenter Finland-funded mass spectrometry instruments at Turku Proteomics Facility were used in this study.

## Ethics declarations

### Author contributions

D.F, P.D. and E.C. contributed to planning, analysis and writing of the manuscript. D.F. and P.J carried out mass-spectrometry-based method development, P.D. carried out sample preparation. P.J. and E.C. supervised the project.

